# Prolonging coagulant activity of factor Xa under hemophilic conditions by site-specific N-glycosylation of the surface-exposed autolysis loop

**DOI:** 10.1101/2021.07.01.450767

**Authors:** Amalie Carnbring Bonde, Jacob Lund, Jens Jacob Hansen, Jakob Rahr Winther, Stefan Zahn, Peter Tiainen, Ole Hvilsted Olsen, Helle Heibroch Petersen, Jais Rose Bjelke

**Affiliations:** Global Research, Novo Nordisk A/S, 2760 Måløv, Denmark; Section for Biomolecular Sciences, Department of Biology, University of Copenhagen, 2200 Copenhagen N, Denmark; Novo Nordisk Foundation Centre for Basic Metabolic Research, University of Copenhagen, 2200 Copenhagen N, Denmark; Research and Development, LEO Pharma, 2750 Ballerup, Denmark

**Keywords:** Coagulation factor X, site-directed mutagenesis, N-linked glycosylation, tissue factor pathway inhibitor, antithrombin, thrombin, hemostasis, bypassing treatment

## Abstract

The regulation of Factor X (FX) is critical to maintain hemostasis. To gain insights to the regulation of the active and zymogen form of coagulation FX, we probed specific molecular interactions by introducing novel N-linked glycosylations on the surface-exposed loop spanning residues 143-150 (chymotrypsin numbering) of FX. Introduction of N-glycans in the autolysis loop of these FX variants decreased Factor VIIa (FVIIa)-mediated activation ~3-fold and prothrombin activation 2- to 10-fold presumably through steric hinderance. Prothrombin activation was, however, recovered in presence of cofactor Factor Va (FVa) despite a reduced prothrombinase assembly. The introduced N-glycans exhibited position-specific effects on the interaction with two FXa inhibitors: tissue factor pathway inhibitor (TFPI) and antithrombin (ATIII). *K_i_* for the inhibition by full-length TFPI of these FXa variants was increased by 7- to 1150-fold, while ATIII inhibition in the presence of the heparin-analogue Fondaparinux was modestly increased by 2- to 15-fold compared to wild type. To probe the *in vitro* hemostatic effect of the FX variants, the thrombin generation potential in FX-depleted plasma was evaluated. When supplemented in zymogen form, the FX variants exhibited reduced thrombin generation activity relative to wild-type FX, whereas enhanced procoagulant activity was measured for activated FX variants with N-glycosylation at positions 148-150. These results indicate that residues of the surface-exposed autolysis loop and residues close by participate in FX activation, proteolytic activity and inhibition of FXa by TFPI and ATIII. In plasma-based assays, a modest decrease in FX-activation rate appeared to compensate for the collective reduction in inhibitor interactions.

## Introduction

Hemostasis is a complex biological system that exquisitely balances between coagulation and fluidity. Upon vascular injury, coagulation is initiated to prevent hemorrhage and involves interrelated protease activation steps and recruitment of thrombocytes, eventually leading to generation of a thrombin burst necessary for efficient fibrin clot formation. A central protease in coagulation is the vitamin K-dependent serine protease FX, which activates prothrombin to thrombin through limited proteolysis, a reaction greatly enhanced by prothrombinase complex assembly with its cofactor, coagulation FVa, on the surface of activated thrombocytes [1, 2]. Following vascular injury, FX is activated by the tissue factor (TF)-FVIIa complex or the factor VIIIa (FVIIIa)-factor IXa (FIXa) complex, where the activation peptide is proteolytically removed to induce maturation of the serine protease domain [3]. FXa is eventually down-regulated by inhibitors to avoid uncontrolled coagulation. Two major inhibitors of FXa are the TFPI [4, 5] and the heparin-dependent serpin ATIII [6, 7].

Reduced activity levels of either cofactor VIII (FVIII) or Factor IX (FIX) lead to the bleeding disorder hemophilia, where the decreased FX-activation and subsequently reduced thrombin generation is expressed phenotypically by prolonged and uncontrolled bleeding. Treatment of hemophilia involves increasing the activation rate of FX by replacing the missing coagulation factors upstream of FX [8] or alternatively using bypassing agents [9, 10]. Novel bypassing concepts aim to slow down the inactivation of FXa by neutralizing the inhibitory activity of ATIII [11] and TFPI [12, 13] both interventions showing hemostatic potential in hemophilia patients.

The autolysis loop, consisting of residues 143-150 (Chymotrypsin numbering), is one of several surface loops that surround the substrate-binding pocket of FXa [14] which is important for substrate and inhibitor specificity and selectivity [15, 16]. Previously, the contributions of the four basic residues in the autolysis loop were evaluated by substituting each residue individually with alanine [15]. The effects of the mutagenesis were only minor on prothrombinase complex assembly, proteolytic cleavage of prothrombin and the inhibition by TFPI.

In the present study, instead of alanine substitutions, we introduced novel N-linked glycosylation motifs into the loop to map surface interactions of FX/FXa. Using targeted N-glycosylation as a tool [16], we probed molecular interaction sites to understand how the loop influences the pro- and anti-coagulant regulation mechanisms and how these mechanisms maintain the balance between coagulation and blood fluidity.

We characterized functional aspects of the selected residues within the autolysis loop with regards to FX zymogen activation, FXa proteolytic activity and anticoagulant regulation of FXa by TFPI and ATIII, and finally the hemostatic potential in FX-depleted plasma.

## Results

### Expression, purification, and characterization of recombinant FX variants

To ensure high-quality production of recombinantly expressed FX glycan variants, especially in terms of the Gla-domain, an anti-Gla immunoaffinity protocol was developed based on a selective FX anti-Gla antibody obtained from an immunization screening in mice. Selectivity of the antibody was shown using Gla-residue substituted variants, in which the antibody was shown to bind only to FX variants with 9 or more γ-carboxylated residues as evaluated using an ELISA experiment (Figure S3). Thus, a combination of immunoaffinity and anion-exchanged chromatography-based purification protocol for producing pure and high-quality recombinant FX variants was established. This method was used for producing wild-type FX and glycan variants hereof selected from a surface-probing library of FX (Figure S1, Figure S2 and Table S1) with N-linked glycosylation motifs introduced in the autolysis loop (at position 143, 147, 149 and 150, respectively (Figure 1A)) masking a surface region spanning up to 50 Å (Figure 1B). A high degree of protein purity, degree of furin processing and homogeneity for all the FX variants were evaluated by reducing SDS-PAGE (Figure 2A) and size-exclusion HPLC analysis (data not shown). From the SDS-PAGE analysis, two bands, heavy chain (HC) and light chain (LC), were observed to exhibit the expected electrophoretic mobility for all the purified FX glycan variants indicating correct furin processing. The heavy chains of the FX variants exhibited decreased electrophoretic mobility relative to wild-type FX consistent with introduction of the intended N-linked glycan (Figure 2A). N-terminal sequencing confirmed correct furin processing and vitamin K-dependent γ-carboxylation of Gla-domain of selected FX variants examined. Together, this showed the effectiveness of the established production protocol for producing high quality recombinant FX variants (Figure S4 and Table S2).

**Figure 1.**
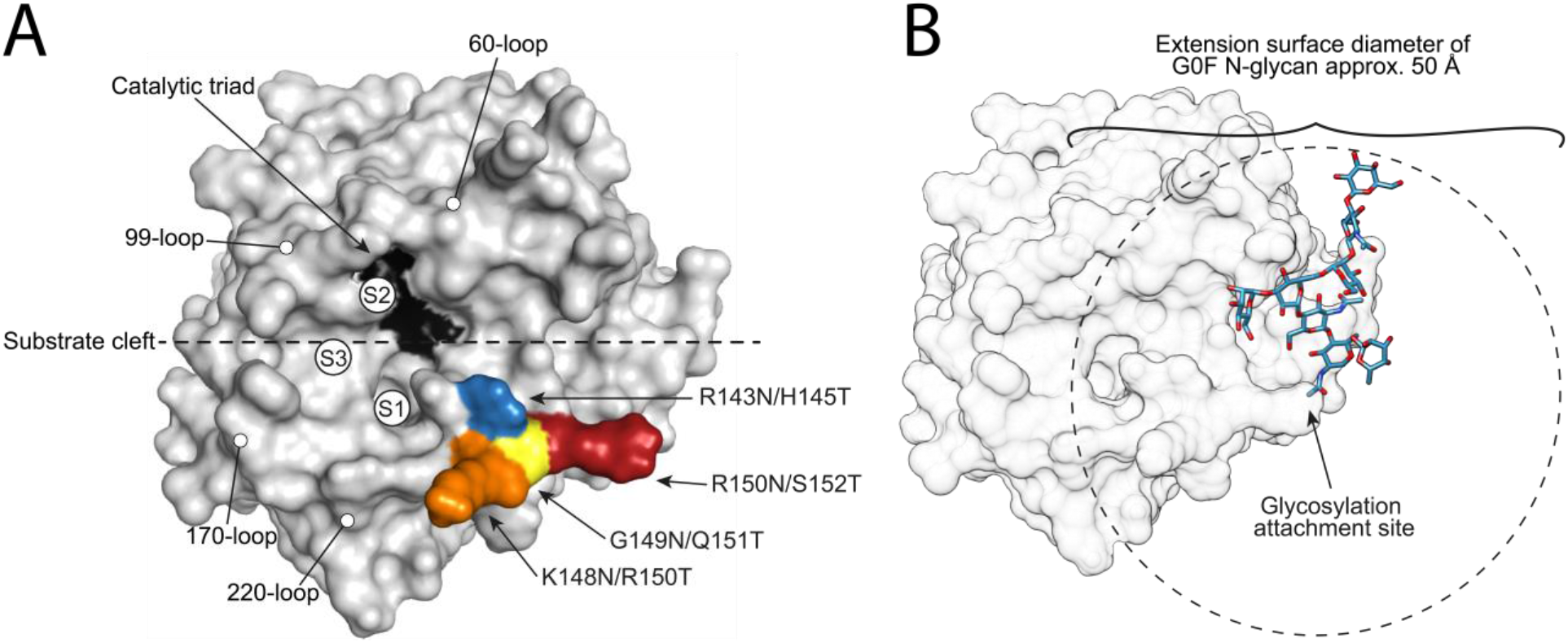
The introduced N-glycosylation sites in the autolysis loop of FX/FXa (PDB ID: 1FAX) and extension surface of N-glycans. (A) Overview of sites selected for introduced N-glycosylation by site-directed mutagenesis. The mutation sites are annotated by chymotrypsin numbering. The active site is indicated in black and selected specificity determining surface loops are indicated. Substrate pockets S1, S2 and S3 are also indicated. (B) Modelling of a representative N-glycan (blue) on position 150 showing the potential mapping surface spanning up to 50 Å in diameter.

**Figure 2.**
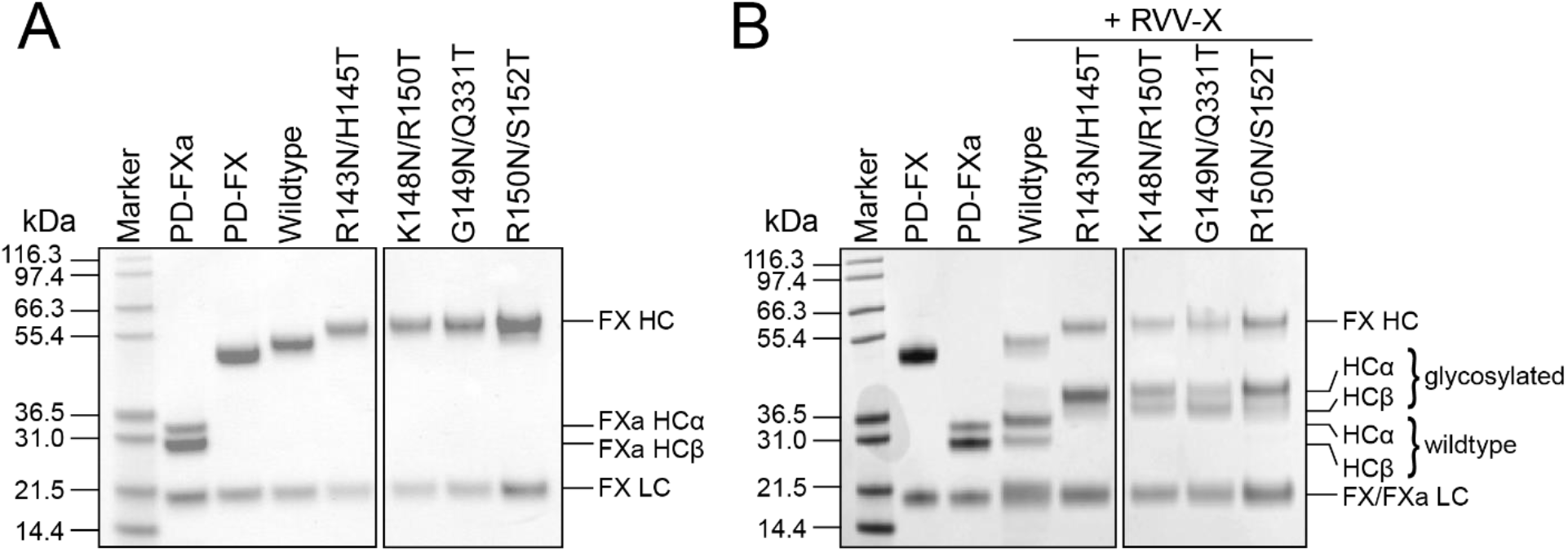
SDS-PAGE analysis of purified FX and FXa variants. (A) Reducing SDS-PAGE analysis of purified zymogen FX variants followed by Coomasie Blue-staining. Plasma-derived (PD) activated FXa and zymogen FX was included for comparison. (B) Reducing SDS-PAGE analysis of RVV-X activated FXa variants followed by Coomasie Blue-staining. PD-FX and PD-FXa were included for comparison.

### Preparative activation of FX variants by RVV-X and active-site probing

To examine the catalytic activity of the recombinant wild-type and mutant FX variants, variants were activated by Russel’s Viper venom X (RVV-X) resulting in the removal of the glycosylated activation peptide of FX by limited proteolysis leading to a decreased mass of heavy-chain. The processed HC was observed as an α-form of approx. 35 kDa and a β-form of approx. 30 kDa for PD-FXa and wild-type FXa, respectively, whereas the light-chain of FXa was observed at approx. 20 kDa for all variants (Figure 2B). The occurrence of two heavy-chain forms is consistent with the additional auto-proteolysis as previously reported [17]. Notably, both cleavage products have been characterized as equally active [18].

From SDS-PAGE analyses, the recombinant FX variants appeared to be only partially activated as residual un-activated HC was observed at approx. 55 kDa for the wild type and approx. 60 kDa for the N-glycan variants (Figure 2B). The partial activation by RVV-X was observed for all variants including wild-type FX, and thus it appeared unlikely to be related to the introduced N-glycan. The Gla-domain has previously been proposed to be an important exosite for RVV-X interaction [19], however the Gla-domain characterization of the variants tested herein demonstrated high level of γ-carboxylation consistent with the employed purification strategy based on a capture anti-Gla immunoaffinity and polish anion-exchange chromatography. As the glycans in the activation peptide have been reported to promote optimal activation of FX [20, 21], the lack of full activation by RVV-X could be ascribed as a possible heterogeneous level of sialylation of the putative N-linked and O-linked glycosylation sites in the activation peptide of FX. This was not investigated further.

Thus, the total active site concentration was determined in all the FXa variant preparations by active-site titration using the tight-binding active-site inhibitor Ecotin. The fraction of activated FXa was determined to be between 50-65% for all variants (Table S1) consistent with the qualitative evaluation by SDS-PAGE (Figure 2B and Figure S2). The concentrations determined by active-site titrations were used in all further evaluations of the FXa variants.

To determine the kinetic parameters for amidolytic activities of the different FXa variants, the hydrolysis of the S-2765 chromogenic FXa peptide substrate was measured. For all variants, except K148N/R150T, the estimated *k_cat_* / *K_m_* values were found to be 2- to 20-fold lower compared to wild-type FXa affecting both *K_m_* and *k_cat_* (Table 1). Interestingly, the estimated *k_cat_* /*K_m_* value for K148N/R150T was two-fold higher than that of wild-type FXa due to a decrease in *K_m_*.

**Table 1.** Active-site probing, activation and proteolytic activity of FX(a) variants.

| Variant | Amidolytic activity |  |  | FFR-cmk inhibition | Activation by extrinsic tenase |  | FXa | Prothrombinase |  |
| --- | --- | --- | --- | --- | --- | --- | --- | --- | --- |
| | $K_m$<br>($\mu\text{M}$) | $k_{cat}$<br>($\text{s}^{-1}$) | $k_{cat}/K_m$<br>($\mu\text{M}^{-1} \text{s}^{-1}$) | $k_{on}$<br>($\mu\text{M}^{-1} \text{s}^{-1} \times 10^3$) | $k_{cat}/K_m^a$<br>($\text{M}^{-1} \text{s}^{-1}$) | $k_{cat}/K_m^b$<br>($\text{M}^{-1} \text{s}^{-1}$) | Initial velocity <sup>c</sup><br>(nM thrombin/min/nM FXa) | Peak velocity <sup>d</sup><br>(nM thrombin/min/nM FXa) | $K_d$<br>(nM) |
| Wild-type | 88 ± 8 | 170 ± 4 | 1.9 | 31.33 ± 1.07 | 13.2 ± 0.3 | 521 ± 10 | 0.214 ± 0.006 | 921 ± 31 | 0.30 ± 0.05 |
| R143N/H145T | 558 ± 56 | 71 ± 2 | 0.1 | 0.32 ± 0.01 | 4.2 ± 0.1 | 202 ± 3 | 0.016 ± 0.001 | 135 ± 4 | 3.01 ± 0.45 |
| K148N/R150T | 29 ± 3 | 114 ± 2 | 3.9 | 151.70 ± 6.77 | 5.0 ± 0.1 | 182 ± 1 | 0.017 ± 0.001 | 824 ± 24 | 1.52 ± 0.17 |
| G149N/Q151T | 219 ± 14 | 99 ± 1 | 0.5 | 3.28 ± 0.12 | 4.2 ± 0.2 | 176 ± 2 | 0.026 ± 0.001 | 1024 ± 14 | 0.82 ± 0.05 |
| R150N/S152T | 81 ± 5 | 93 ± 2 | 1.1 | 23.74 ± 0.44 | 4.8 ± 0.5 | 176 ± 3 | 0.098 ± 0.002 | 881 ± 19 | 0.58 ± 0.06 |

<sup>a</sup> FX activation by 200 nM FVIIa in solution.<sup>b</sup> FX activation by 20 nM FVIIa and 200 nM sTF.<sup>c</sup> The initial velocities of thrombin activation by 20 nM FXa in presence of 50 $\mu\text{M}$ 25:75 PS:PC.<sup>d</sup> The peak of thrombin activation by prothrombinase in presence of 10 pM FXa and 50 $\mu\text{M}$ 25:75 PS:PC.**Table 2. Kinetic parameters for inhibition of FXa variants by TFPI and ATIII.**

To further investigate effects of the introduced N-glycans on the FXa active site, the substrate binding pocket was probed with H-D-Phe-Phe-Arg-chloromethylketone (FFR-cmk) inhibitor. For all FXa variants, binding of FFR-cmk was perturbed to a similar degree as observed for the amidolytic activity profiling (Table 1).

### Activation of FX variants by extrinsic (TF-FVIIa) and intrinsic (FVIIIa-FIXa) tenase

The effects of introducing N-linked glycosylation sites into the FX autolysis loop on the activation by FVIIa was evaluated in absence or presence of soluble Tissue-Factor (sTF). The *k_cat_* / *K_m_* for FVIIa activation of the autolysis loop variants were reduced 3- to 4-fold compared to wild-type FX (Table 1). Similar reductions were observed with FVIIa/sTF indicating that the N-glycan had no effect on the interaction of zymogen FX with sTF in complex with FVIIa.

No difference was observed between wild type and the N-glycan FX variants when the activation by FIXa/FVIIIa complex was evaluated (Figure S5).

### Proteolytic activity of FXa variants

To measure the ability of FXa variants to activate its natural substrate, prothrombin, the proteolytic activities were determined on a phospholipid surface in absence or presence of FVa. In absence of FVa, the proteolytic activity of R150N/S152T was reduced 2-fold compared to wild type, while the activities of the remaining mutants were reduced 8- to 13-fold (Figure 3A and Table 1). As K148N/R150T displayed increased amidolytic activity compared to wild-type FXa, the reduced proteolytic activity indicated that the introduced N-glycan interferes with prothrombin binding.

**Figure 3.**
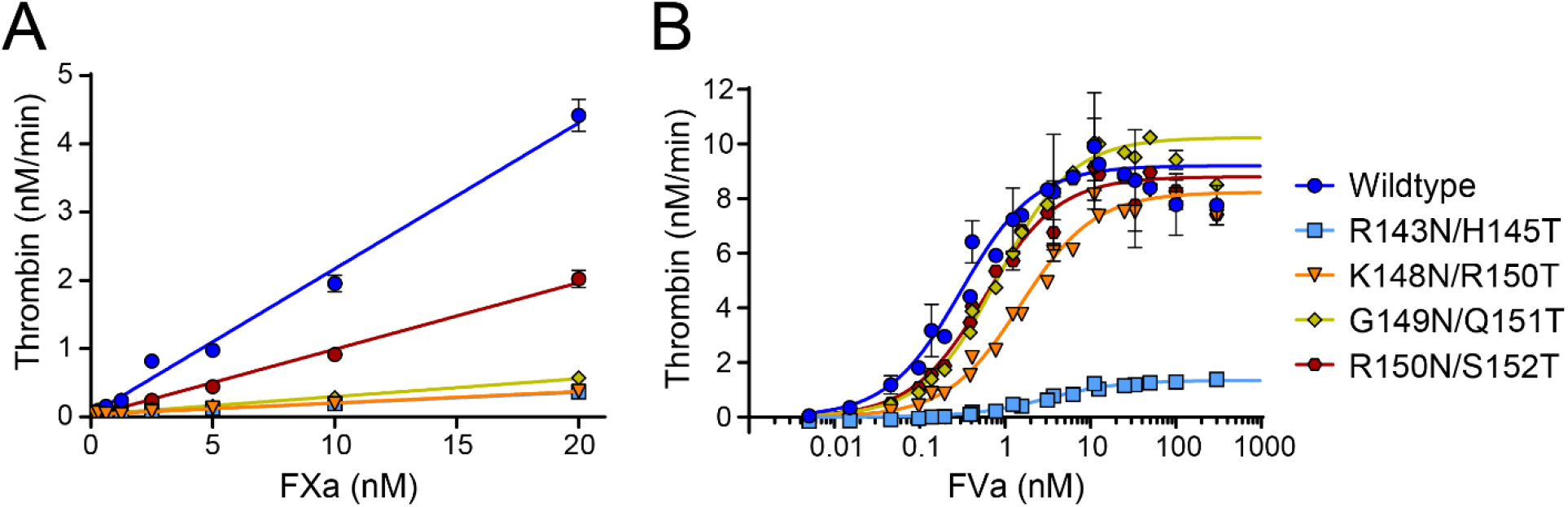
Proteolytic activity and prothrombinase assembly of FXa variants. (A) Thrombin activation by 0-20 nM FXa in presence of 50 μM 25:75 PS:PC. (B) The effect of FVa on thrombin activation by 10 pM FXa in presence of 50 μM 25:75 PS:PC. Thrombin activity rates obtained at high FVa concentrations with a distinctive hook effect were not included in the fit.

However, in presence of FVa, the proteolytic activity was essentially restored for all variants except R143N/H145T even though the apparent prothrombinase assembly affinities were decreased for all variants. The *K_d_* was increased with 2- to 3-fold for G149N/Q151T and R150N/S152T, 5-fold increase for K148N/R150T, while a 10-fold increase was observed for R143N/H145T (Figure 3B and Table 1). As presence of FVa did not restore the proteolytic activity of R143N/H145T, this, in agreement with amidolytic activity and FFR-cmk inhibition, indicated a perturbed active-site even in the presence of FVa.

### Inhibition by TFPI and ATIII

The effects of introducing N-linked glycosylation sites into the FX autolysis loop on the inhibition by the isolated Kunitz domain 2 of TFPI (TFPI-K2) and full-length TFPI was studied by following progress curves of chromogenic substrate hydrolysis at different concentrations of described TFPI variants (Figure 4 and Table 2). The *K_i_* for TFPI-K2 and full-length TFPI inhibition of wild-type FXa were determined to be 15 nM and 0.06 nM (Table 2), respectively, which is in agreement with previous reports on TFPI inhibition of FXa [22–25]. For TFPI-K2 inhibition, the variant G149N/Q151T was inhibited similarly to wild type, while a moderate 2-fold increase of *K_i_* was observed for the variant R150N/S152T. No inhibition of R143N/H145T and K148N/R150T were observed under the tested conditions. For full-length TFPI, the introduced N-glycan appeared to hinder the interaction between TFPI and the FXa active-site of the variants as evident by 1150-, 17-, 33- and 7-fold increase in *K_i_* values for R143N/H145T, K148N/R150T, G149N/Q151T and R150N/S152T, respectively (Table 2). Interestingly, the variants R143N/H145T and K148N/R150T, which did not show inhibition by TFPI-K2, were inhibited by full-length TFPI, although with increased *K_i_* values.

**Table 2.** Kinetic parameters for inhibition of FXa variants by TFPI and ATIII.

| Variant | TFPI-K2 | Full-length TFPI | ATIII | ATIII-H5 |  |
| --- | --- | --- | --- | --- | --- |
| | $K_i$<br>(nM) | $K_i$<br>(nM) | $K_i$<br>(nM) | $K_i$<br>(nM) | $K_{i,uncat}/K_{i,H5}$ |
| Wild-type | 14.5 ± 0.2 | 0.06 ± 0.01 | 699 ± 52 | 4.5 ± 0.5 | 156 |
| R143N/H145T | *N.D.*<sup>a</sup> | 68.71 ± 17.54 | *N.D.*<sup>a</sup> | *N.D.*<sup>a</sup> | *N.D.*<sup>a</sup> |
| K148N/R150T | *N.D.*<sup>a</sup> | 1.19 ± 0.28 | 216 ± 36 | 18.6 ± 6.2 | 11 |
| G149N/Q151T | 28.6 ± 0.83 | 1.85 ± 0.35 | 1924 ± 921 | 69.9 ± 23.4 | 27 |
| R150N/S152T | 26.2 ± 0.85 | 0.40 ± 0.09 | 691 ± 220 | 9.1 ± 4.4 | 77 |

<sup>a</sup> N.D.; not determinable, as the variant was not inhibited at the assay conditions

**Figure 4.**
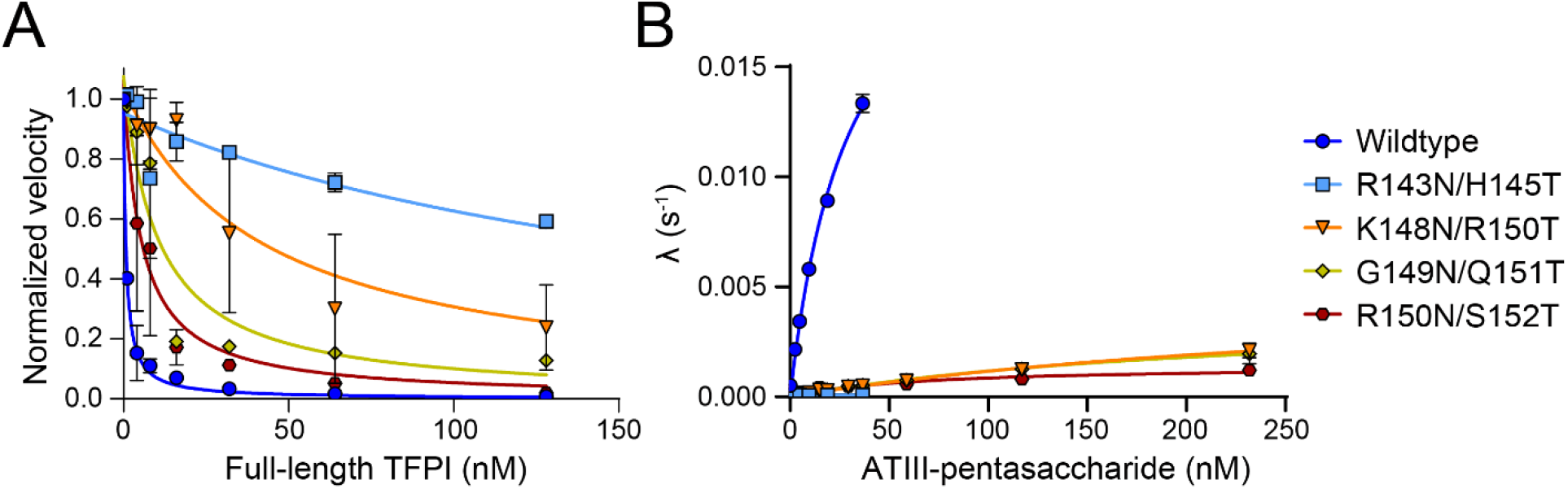
Inhibition of FXa by TFPI and ATIII. (A) Determination of the dissociation constant, ***K_i_***, for inhibition of FXa by full-length TFPI. The steady state velocities were fitted from the progress curves of FXa hydrolysis of S-2765 in presence 1-128 nM TFPI. Steady state velocities were replotted as a function of TFPI concentration. (B) Dependence of the observed decay rate, ***λ*** (***s***^−**1**^), on ATIII-pentasaccharide concentrations. The decay rate, ***λ***(***s***^−**1**^), were measured for reactions of FXa hydrolysis of 500 μM S-2765 in presence of (A) 0.125 – 4 μM ATIII or (B) 1000 nM ATII and 3.125 – 250 nM pentasaccharide. Loss of activity curves were fitted to single exponential decay function and decay rates, ***λ***(***s***^−**1**^) and were replotted as a function of ATIII-pentasaccharide concentrations.

The decay rates for FXa inhibition by ATIII in absence and presence of the heparin-related pentasaccharide, Fondaparinux (H5), were determined under pseudo first-order conditions (Figure 4). In absence of pentasaccharide, only minor effects were observed on the inhibition constant, *K_i_*, for all variants except R143N/H145T, which was not inhibited under the tested conditions. In presence of pentasaccharide, the inhibition constant was increased by 2-fold for R150N/S152T, and 4- to 16-fold for K148N/R150T and G149N/Q151T (Table 2).

### Effect of FXa variants on thrombin generation in plasma

Finally, the hemostatic potential of FX variants in plasma was evaluated in a thrombin generation assay using FX-deficient plasma (Figure 5A). Surprisingly, the thrombin peak was clearly reduced for all N-glycan variants compared to wild type, with a 4-fold decrease for variants G149N/Q151T and R150N/S152T, while variant K148N/R150T was reduced 50-fold. Variant R143N/H145T was not tested due to the poor proteolytic activity observed for this variant.

**Figure 5.**
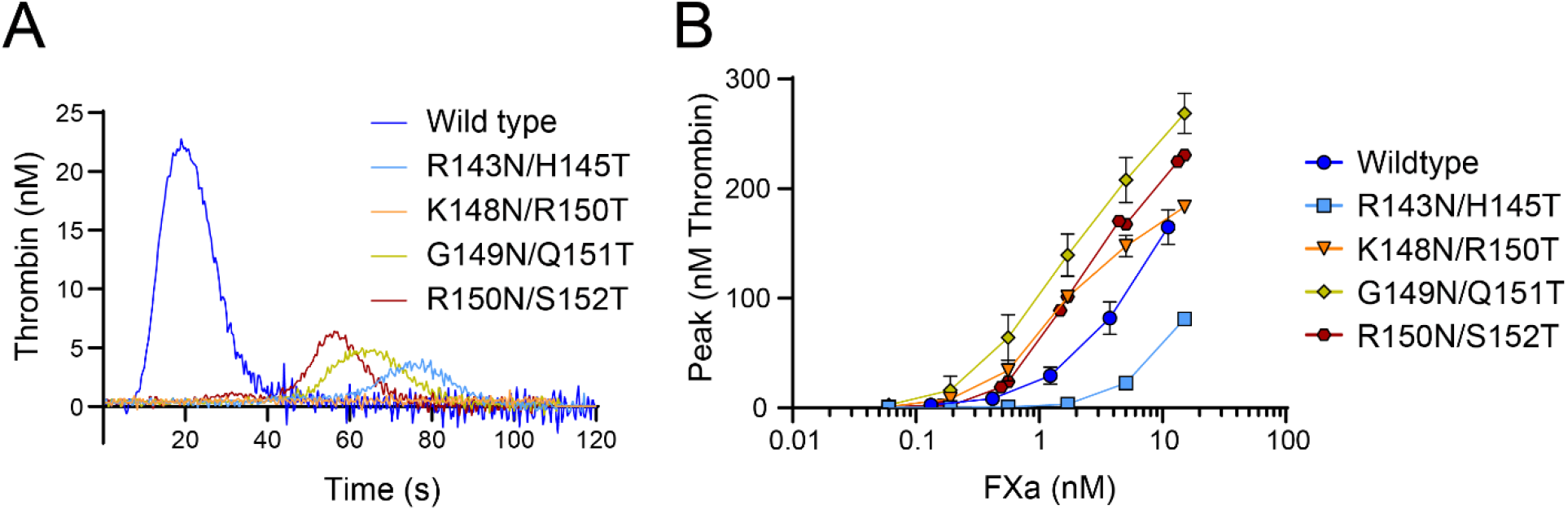
Thrombin generation by FX/FXa variants in Thrombin generation assay under hemophilic conditions. Thrombin generation was measured in FX-depleted platelet-poor plasma supplemented with neutralizing FVIII antibodies. (A) Plasma was supplemented with 136 nM zymogen FX variants and triggered by 1 pM TF. (B) Thrombin generation was initiated by addition of 0-15 nM FXa variant.

To eliminate the activation step, we instead initiated the thrombin generation by addition of the activated variants (Figure 5B and Figure S6). Under these conditions, G149N/Q151T, K148N/R150T and R150N/S152T all exhibited increased potency as compared to wild-type FXa, whereas R143N/H145T displayed compromised thrombin generation potential.

As the FXa preparations contained a fraction of zymogen FX, we evaluated whether this could affect the assay by titrating increasing concentrations of plasma-derived FX in the Thrombin generation assay triggered with 0.2, 1 or 5 nM plasma-derived FXa, however no effect was observed (Figure S7).

## Discussion

In this study, we have examined the hemostatic potential of FX(a) variants with N-glycans introduced on the surface of the protease domain of FX. As modelled in Figure 1B, introduction of an N-glycan on the surface introduce a spatial shield spanning up to 50 Å in diameter representing a powerful tool for surface mapping. Thus, four variants with N-glycans introduced to the autolysis loop (residue 143-150 in chymotrypsin numbering) displayed interesting biochemical properties of the autolysis loop and surface vicinity. These FXa variants emerged with improved thrombin generation capability obtained by protecting the pivotal coagulation factor against inhibition. Interestingly, while located within close structural proximity in FXa, the individual N-linked glycans appeared to exert different mechanism of actions in relation to procoagulant activity and inhibition of the FXa-variants.

We observed impaired amidolytic activities and decreased second-order rate constant for inhibition by the peptidyl inhibitor FFR-cmk for R143N/H145T, G149N/Q151T, and R150N/S152T, indicating that the mutagenesis had affected the active-site structure or reactivity of the catalytic pocket of these variants. For these variants, the reduced active-site inhibition correlated well with both the determined amidolytic activity (against S-2765) and proteolytic activity (against prothrombin) indicating that the introduced N-glycosylations affected the active site and not the interaction with prothrombin. In contrast, K148N/R150T displayed improved amidolytic activity compared to wild type, but reduced proteolytic activity, hence we speculate that this is due to steric hinderance of prothrombin binding by the introduced N-glycosylation at position 148. Interestingly, in presence of phospholipids and FVa, all variants except R143N/H145T, activated prothrombin similar as wild-type FXa. If the consequence of inserting an N-glycosylation at R143N/H145T, G149N/N151Q and R150N/S152T was an altered active-site structure or reactivity of the active site to a less favourable conformation, presence of FVa only seemed to nullify this effect for variant G149N/N151Q and R150N/S152T. Furthermore, FVa appeared to cancel out the possible hinderance between FXa and prothrombin introduced by the N-glycan at K148N/R150T, indicating that the cofactor either was able to push the glycan out of the way or was inducing a conformational change in this region, hence eliminating the effect of N-glycan at this position. It has been reported previously [26, 27] that prothrombin activation can occur via two mechanistically different pathways, comprising intermediates prethrombin or meizothrombin, depending on absence or presence of phospholipids and cofactor.

An acidic patch on the surface of TFPI-K2 have been proposed to interact with the basic residues of the autolysis loop of FXa [28]. Nevertheless, only a minor effect was observed in a mutagenesis study of K148A, while R143A and R150A was inhibited normally by full-length TFPI [15]. In this study, the insertion of an N-glycosylation at G149N/Q151T and R150N/S152T was more critical for the interaction between FXa with full-length TFPI compared to inhibition by Kunitz 2 domain alone. This was evident from the 33- and 7-fold increase in *K_i_* values for G149N/Q151T and R150N/S152T, respectively, while inhibition of G149N/Q151T by TFPI-K2 was not affected and only a 2-fold increase in *K_i_* was observed for R150N/S152T. This indicated that the N-glycosylations at position 149 and 150 interacted with surface sites of TFPI outside Kunitz domain 2. On the other hand, TFPI-K2 inhibition of the variants R143N/H145T and K148N/R150T was completely disrupted. Full-length TFPI was able to alleviate the effect of the inserted N-glycosylation at position 143 and 148, although the *K_i_* values were still increased by 1150- and 17-fold, respectively. In conclusion, full-length TFPI exosite interactions on FXa rely less on interaction with the autolysis loop and can execute inhibition, even though interaction between Kunitz domain 2 and FXa active site is perturbed.

A previous study showed that substituting R143 or K148 of FXa for alanine residues slightly improved ATIII inhibition in the absence of cofactor, suggesting that charge and/or size of these side chains were restricting inhibition by the serpin [15]. Hence introduction of a bulky N-glycosylation moiety was expected to have the opposite effect from introduction of an alanine residue. Indeed, R143N/H145T was not inhibited by ATIII under the measured conditions, neither in absence nor presence of pentasacharide H5. For the remaining three FXa-variants tested, the rate constants determined were all reduced both in the presence and absence of pentasacharide as compared to wild-type FXa. However, due to the effect of the introduced N-glycans on the *K_m_* for S-2765 proteolysis, the resulting *K_i_* for ATIII inhibition in the absence of oligosaccharide-boosting were only slightly affected. In contrast, *K_i_* for the inhibition of the FXa-variants by ATIII boosted by the pentasacharide H5 were significantly increased compared to wild-type FXa. Previously, the pentasacharide H5 were shown to significantly decrease the *K_i_* for ATIII-mediated inhibition of recombinant, wild-type FXa [29] presumably by inducing conformational changes in ATIII promoting FXa inhibition as shown for heparin based oligosaccharides [30, 31]. Hence, our data suggests that the introduced N-glycans render FXa less sensitive to the imposed conformational changes to ATIII upon binding heparin or derived oligosaccharides. Noticeably, R150 was previously found to interact productively with ATIII, in particular in presence of pentassacharide [15]. It is possible that the effect of K148N/R150T on ATIII inhibition in presence of pentassaccharide cannot be ascribed solely to the introduction of an N-glycosylation at position 148, as the mutation R150T could have an effect by itself. Interestingly, only minor effects were observed for ATIII inhibition of R150N/S152T in presence of pentassacharide, as the variant was only impaired 2-fold in contrast to the data reported for the equivalent R150A mutant. Of all four autolysis loop variants, the mutant G149N/Q151T affected ATIII inhibition the most.

Although we observed reduced activation rates by TF-FVIIa of the FX autolysis loop N-glycan variants, we hypothesized that the variants K148N/R150T, G149N/Q151T and R150N/S152T could improve the thrombin generation peak in hemophilia A plasma as they showed wild-type like proteolytic activity and were better protected against inhibition by TFPI and ATIII compared to wild type. However, surprisingly, this was not the case as the N-glycan variants were 4- to 50-fold less effective in generating Thrombin peak in hemophilia A plasma as compared to wild-type FX. Furthermore, by instead pre-activating the FX variants prior to supplementation to hemophilia A-like plasma, the three out of four N-glycan variants exhibited superior potency compared to wild-type FXa. Hence these results indicate the importance of the initial activation steps of coagulation where the extrinsic tenase complex activates initial amounts of FX, as small perturbations of the FX activation rate outweigh significant abrogation of inhibitor interactions. This suggest that introduction of site-specific N-glycans or similar moieties on the surface of FX, such as the autolysis loop, can render enhanced inhibitor protection, which could be exploited therapeutically. Of notice to this is a reporting that FX Ala mutants of the autolysis loop display decreased endothelial barrier protection via modulated specificity towards protease-activated receptor-2 [32].

Specificity determination driven via the autolysis loop have also been reported via chimeric loop swaps between FX, protein-C and trypsin emphasizing the importance of the loop among enzymes of the coagulation cascade and other pathways [33].

In conclusion, we observed distinct biochemical effects for FX N-glycan variants designed to probe the autolysis loop and structural vicinity of FX for impact on various phases of the coagulation cascade. Further, introduction of novel N-glycan side chains to the molecular surface of coagulation factors and other proteins can effectively map large surface areas.

## Experimental Procedures

### Reagents

Plasma coagulation factors and 25:75 PS:PC (phosphatidylserine-phosphatidylcholine) phospholipids were purchased from Haematologic Technologies Incorporated (Vermont, USA). Chromogenic substrates, S-2765 and S-2238, were purchased from Instrumentation Laboratory (Massachusetts, USA). RVV-X was purchased from Sekisui Diagnostics (Massachusetts, USA). Ecotin was purchased from Molecular Innovations (Michigan, USA). Phe-Phe-Arg-chloromethylketone (FFR-cmk) was purchased from Bachem (Switzerland). Recombinant soluble tissue factor (sTF), recombinant FVIIa, and recombinant TFPI variants were produced and provided by Novo Nordisk A/S. All enzymatic assays were carried out in a reaction buffer consisting of 50 mM Hepes, 100 mM NaCl, 10 mM CaCl_2_, 0.1 % PEG8000, 1 mg/ml BSA adjusted to pH 7.4.

### Mutagenesis, transfection and expression of recombinant factor X variants

Chimeric FX cDNA vectors encoding full-length, mature factor X (FX) protein preceded by the prothrombin signal sequence and propeptide were purchased from GenArt (Thermo Fisher Scientific). N-glycosylation acceptor sites were introduced successfully at modelled surface exposed sites of publicly available crystal structure of FXa (PDB id 1FAX and residues 60, 61, 74, 93, 96, 143, 146, 147, 149, 150, 173, 185, 188, 222 and 243) using the motif Asn-X-Thr. FX variants were expressed in CHO cells using QMCF expression system (Icosagen Cell Factory, Tartumaa, Estonia) according to manufacturer’s instructions.

### Anti-Gla antibody and purification of factor X

To produce a FX anti-Gla antibody for a purification protocol, RBF mice were immunized with human FX emulsified in Freuds complete adjuvants for the first injection followed by two additional biweekly injection with incomplete Freuds adjuvants. Sera were tested in a direct FX ELISA and the mouse with the highest antibody titer was used in a fusion. After an i.v. booster injection at day −3 spleenocytes were isolated and fused to FOX-NY myeloma cells using electrofusion. Fused cells were seeded in 96 well plates and supernatant from hybridoma clones were screened after 10 days cultivation at 37°C in an indirect ELISA for binding to FX and to Gla-domain deleted FX. Only clones with FX binding which showed no binding to Gla domain deleted FX were selected resulting in the clone m-FX-10F2A2, which also bound full length FX in a calcium-dependent way. The DNA sequences for the VL and VH of FX-10F2A2 were cloned and CHOK1SV GKSO cell line was established based on an electroporation protocol and screening for high-titer antibody-producing cells using a ForteBIO instrument. The selected clone was cultivated in a 15-L bioreactor using a fed batch cultivation protocol producing gram-scale of the anti-Gla antibody, designated 10F2A2. The monoclonal antibody was purified using a Protein-A based purification protocol setup on an Äkta Explorer (Cytiva). Using the purified anti-Gla antibody, an immunoaffinity resin was developed by coupling the anti-Gla antibody to CNBr-Sephraose-4 Fast Flow resin (Cytiva). Capacity of selective binding of FX variants with Gla-residue substitutions were performed based on an optimized purification protocol. The FX Gla-residue mutants tested for the Gla-domain selectivity were these: FX-E^39^D, FX-E^32/39^D, FX-E^29/32/39^D, FX-E^26/29/32/39^D, FX-E^25/26/29/32/39^D, FX-E^20/25/26/29/32/39^D, FX-E^19/20/25/26/29/32/39^D, FX-E^16/19/20/25/26/29/32/39^D, FX-E^14/16/19/20/25/26/29/32/39^D, FX-E^7/14/16/19/20/25/26/29/32/39^D, FX-E^6/7/14/16/19/20/25/26/29/32/39^D.

The cloned murine anti-FX Gla antibody VL/VH sequences are shown hereunder:

VH:

QVQLKQSGPGLVQPSQSLSITCTVSGFSLTT YGVHWVRQSPGDGLEWLGVIWSSGSTDY YAPFKSRLTISKDNSKSQVFFKMNSLQAQD TAIYYCATLGGFAYWGQGTLVTVSA

VL:

QIVLTQSPAIMSASPGEKVTLTCQASSSVSS SYFYWYQQKPGSSPKLWIKSTSKLASGVPA RFSGSGSGTSYSLTISSMEAEDAASYFCHQ WSSYPLTFGAGTKLELI

The developed protocol was used for purifying expressed FX glycan variants from cell supernatants setup as a two-step integrated chromatographic approach on an Äkta Xpress instrument (Cytiva). Prior to the first purification step, cell supernatants were supplemented with 10 mM CaCl_2_. To decrease proteolytic degradation, 5 mM benzamidine was added to cell supernatants and pH was adjusted to 6.0. The anti-Gla antibody coupled resin was packed in a Tricorn 10/50 column and bound FX protein applied to the column was eluted in a buffer containing 60 mM EDTA. Subsequently, the FX variants were further purified by anion exchange chromatography to yield fully γ-carboxylated Gla-domain using Source 15Q resin packed in a Tricorn 5/20 column (Cytiva). Following immediate application of eluate from the immunoaffinity chromatography step onto the Source 15Q column, this column was washed with 300 mM ammonium acetate prior to elution of the FX protein using 10 mM Histidine, pH 6.0, 100 mM, 20 mM CaCl_2_.

### Activation of factor X variants by RVV-X

FX variants were converted to active forms by incubation with RVV-X enzyme coupled to CNBr-activated Sepharose 4 Fast Flow resin (Cytiva) at 37°C for 24 h in 10 mM histidine, 20 mM CaCl_2_, 100 mM NaCl, pH 6.0. The ratio of RVV-X to FX was 1:15. The activation reaction was filtered through a 4 mm syringe filter 0.45 μm PVDF (Chromacol, Greyhound Chromatography) in order to separate FXa from coupled RVV-X.

### Active-site concentration determinations of factor Xa variants

Active-site concentrations of the recombinant FXa variants were determined by titration with the active site inhibitor ecotin assuming a 1:1 stoichiometry. Initially, the concentrations of FXa variants were based on the measured absorbance of the stock solution at 280 nm. The fraction of active-sites was estimated by dividing the active-site concentration with concentration measured by absorbance. The determined active site concentrations were used for all assays used to analyze FXa variants.

### Kinetics of chromogenic substrate hydrolysis

The initial rates of hydrolysis of various concentrations of the chromogenic substrate S-2765 by 2 nM FXa variants were fitted to the Michaelis-Menten equation to derive a *k_cat_* and *K_m_*.

### Inhibition by active-site inhibitor FFR-cmk

The rate of inactivation of FXa variants by FFR-cmk was measured in a discontinuous assay under pseudo first-order rate conditions. At different timepoints (0-60 min), 1 nM FXa variant was added to a reaction mixture containing FFR-cmk (0.125-1 μM) in 50 mM Hepes, 100 mM NaCl, 5 mM CaCl_2_, 0.1% PEG8000, 1 mg/ml BSA, pH 7.4 at room temperature. The remaining enzyme activity as a function of time was measured after the addition of 0.5 mM S-2765 and the observed initial rates of S-2765 hydrolysis were fitted to exponential decay function to derive the observed decay rate, *λ*. From the plot of observed decay rate as a function of FFR-cmk showed a straight line, thus the rate constants *k_on_* and *k_off_* were determined according:

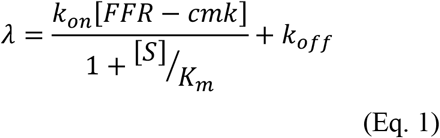

### Factor X activation by FVIIa in solution

The activation of FX variants by either 200 nM FVIIa or 20 nM FVIIa pre-incubated with 200 nM sTF were measured following incubation with 0-500 nM FX variant for 20 min. Reactions were quenched by addition of excess EDTA and initial rates of FXa generation were determined in the presence of 0.5 mM S-2765 by relating the measured amidolytic activities to a standard curve with known concentrations of FXa variants. A *k_cat_* / *K_m_* was derived by fitting to linear regression.

### Factor X activation by FIXa/FVIIIa

For the activation of FX by FIXa/FVIIIa complex, 2 nM FVIII was initially incubated with 0.5 mM PS:PC and 20 nM thrombin for 1 min, followed by addition of 240 nM Hirudin in order to quench thrombin activity. Immediately, 2 nM FIXa was added to the reaction and mixed for 20 seconds prior to addition of 100 nM FX. Activation of FX by FIXa/FVIIIa was quenched by addition of excess EDTA after 20 seconds and initial rates of FXa generation were determined in the presence of 0.5 mM S-2765 by relating the measured amidolytic activities to a standard curve with known concentrations of FXa variants.

### FXa-mediated activation of prothrombin

The activation of prothrombin by FXa variants were measured in presence of 50 μM 25:75 PS:PC vesicles. Following incubation of 1 μM prothrombin with 0-20 nM FXa for 3 min, reactions were quenched by addition of excess EDTA and 50 nM ecotin. The initial rates of thrombin generation were determined in presence of 0.5 mM S-2238 by relating the measured amidolytic activities to a standard curve with known concentrations of α-thrombin.

### Apparent dissociation constant K_d(app)_ for FVa

The apparent dissociation constant, *K_d(app)_*, for FVa interaction with FXa variants were determined from the effect of varying concentrations of FVa on the prothrombin activation rate in presence of phospholipid surface. 10 pM FXa was incubated with 0-300 nM human FVa and 50 μM 25:75 PS:PC vesicles for 5 min. The activation reaction was initiated by addition of 1 μM human prothrombin and quenched after 5 min with excess EDTA. Thrombin generation was evaluated from initial rates of S-2238 hydrolysis and related to the concentration of α-thrombin from a standard curve with known concentrations of thrombin. The dissociation constant *K_d(app)_* for interaction of FXa variants with human FVa were extracted from the titration curves by fitting the data by nonlinear least squares regression analysis to the quadratic binding equation as described in [34]:

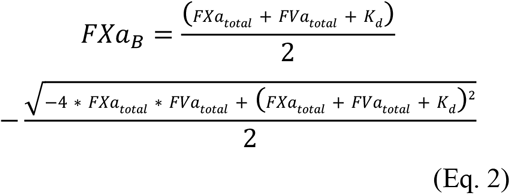

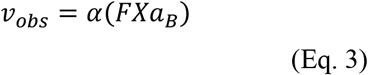

*FXa_total_* and *FVa_total_* refer to total concentrations of the two reactants, *K_d_* is the apparent dissociation constant for the interaction between FXa and FVa. The observed initial rate of S-2238 hydrolysis *v_obs_* is related to the concentrations of FXa bound to FVa by the term *α* (observed initial rate with 10 pM FXa completely saturated with FVa).

### Inhibition of FXa by TFPI

The inhibition of FXa variants by TFPI-K2 and full-length TFPI recombinant variants was determined as previously described in [25]. Various concentrations of TFPI variants was incubated with S-2765 for 5 min (0.5 mM for reactions with TFPI-K2 and 1 mM for reactions with full-length TFPI. Subsequently, 0.4 nM FXa was added and hydrolysis of S-2765 was monitored continuously for 60 min. The data obtained from the progress curves was either fitted by linear regression (TFPI-K2) from which the slope *v_s_*, was obtained, or to the integrated rate equation for slow-tight binding inhibition (full-length TFPI) as described in [35]:

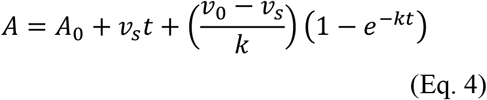

Where *A* is the absorbance at time *t*, *A*_0_ is the absorbance at time 0, *v*_0_ is the initial velocity, *v_s_* is the velocity at steady state, and *k* is the rate of inhibition. The overall dissociation constant *K_i_* was determined from the plot of *v_s_* as a function of TFPI concentration using [35]:

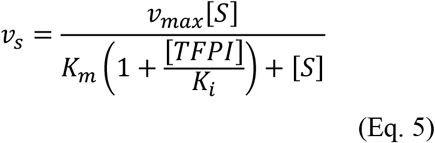

Where *v_max_* is the value of *v_s_* in absence of TFPI, [*S*] is the concentration of S-2765, *K_m_* is the Michaelis-Menten constant for FXa variant towards S-2765.

### Inhibition of FXa by ATIII

The rate of inactivation of FXa variants by ATIII in both absence and presence of the pentasaccharide Fondaparinux was measured under pseudo first-order rate conditions by a discontinuous assay method as described in [36, 37]. In absence of H5, 1 nM FXa was incubated for 0 – 120 min with 0.125 – 4 μM ATIII. In presence of H5 0.4 nM wild-type FXa was incubated for 0 – 20 min with 200 nM ATIII and 3.125 – 50 nM H5. For FXa variants, 1 nM FXa was incubated for 0 – 60 min with 1000 nM ATIII and 15.625 – 250 nM H5. The remaining enzyme activity as a function of time was measured after the addition of 0.5 mM S-2765 and 0.5 mg/ml polybrene (only for reactions with H5) and the observed initial rates of S-2765 hydrolysis were fitted to exponential decay function to derive the observed decay rate, *λ*. From the plot of observed decay rate as a function of ATIII or ATIII-pentasaccharide complex concentration, hyperbolic curves were obtained and fitted to equation 4 or 5, as described in [36]:

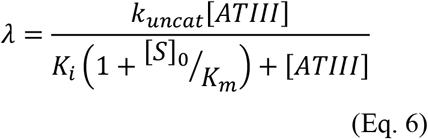

or in presence of H5,

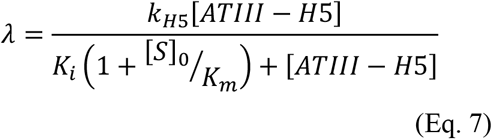

Where *k_uncat_* and *k*_*H*5_ represents the first-order rate constant for conversion of intermediate FXa-ATIII complex to a stable complex, *K_i_* is the dissociation constant for binding of FXa to ATIII, [*S*]_0_ is the concentration of S-2765 and *K_m_* is the Michaelis-Menten constant for S-2765 hydrolysis by the protease. The apparent second-order rate constant *k*_2_ is represented by the initial slope and is calculated from the ratio of ^*k*^/*k_i_*.

For ATIII inhibition in presence of H5, complex concentrations were calculated from the dissociation constant *K_D_* for the interaction between ATIII and H5 and the total concentrations using the quadratic equation [36]. *K_D_* for Fondaparinux was determined to 60 nM in [6].

### Thrombin generation assay

The effect of the FX/FXa N-glycan variants on thrombin generation was measured in FX-depleted plasma (Affinity Biologicals, Ontario, Canada) supplemented with neutralizing FVIII antibodies. For testing zymogen variants, FX-depleted plasma was supplemented 136 nM zymogen FX variant and was triggered by TF (1 pM, PPP-low). For evaluation of activated variants, coagulation was initiated by addition 0-136 nM activated FXa variant and triggered by recalcification. Thrombin activity was assessed by calibrated automated thrombin generation measurements [38].

## Supporting Information

This article contains supporting information.

## Conflicts of interest

The authors declare that they have no conflicts of interest with the contents of this article. A.C.B, J.L, J.J.H, S.Z, P.T and J.R.B are currently Novo Nordisk employees (Novo Allé, DK-2880 Bagsvaerd, Denmark).

## Abbreviations

FX: factor X
FVIIa: Factor VIIa
sTF: soluble tissue factor
TFPI: tissue factor pathway inhibitor
ATIII: antithrombin
FVIII: cofactor VIII
FIX: Factor IX
HC: heavy chain
LC: light chain
RVV-X: Russel’s Viper venom X
FFR-cmk: H-D-Phe-Phe-Arg-chloromethylketone
FVa: cofactor Va
K2: kunitz domain 2
TGT: thrombin generation test
PS:PC: phosphatidylserine-phosphatidylcholine
Gla: γ-carboxylated glutamic acid
H5: pentasaccharide Fondaparinux

